# Nucleic acid receptor ligands improve vaccination efficacy against the filarial nematode *Litomosoides sigmodontis*

**DOI:** 10.1101/2022.07.11.499516

**Authors:** Johanna F. Scheunemann, Frederic Risch, Julia J. Reichwald, Benjamin Lenz, Anna-Lena Neumann, Stephan Garbe, Stefan J. Frohberger, Marianne Koschel, Jesuthas Ajendra, Maximilian Rothe, Eicke Latz, Christoph Coch, Gunther Hartmann, Beatrix Schumak, Achim Hoerauf, Marc P. Hübner

**Affiliations:** Institute for Medical Microbiology, Immunology and Parasitology, University Hospital Bonn, Bonn, Germany; Clinic for Radiotherapy and Radiation Oncology, University Hospital Bonn, Bonn, Germany; Institute for Innate Immunity, University Hospital Bonn, Bonn, Germany; Institute of Clinical Chemistry and Clinical Pharmacology, University Hospital Bonn, Bonn, Germany; Nextevidence GmbH, Munich, Germany; German Center for Infection Research (DZIF), Partner Site Bonn-Cologne, Bonn, Germany

**Keywords:** filariae, *Litomosoides sigmodontis*, 3pRNA, poly(I:C), type I interferon, vaccination, nucleic acid receptor, helminth

## Abstract

Infections with helminths affect more than one billion people worldwide. Despite an urgent need there is no vaccine available that would confer long lasting protection against helminth infections. Previous studies indicated that a vaccination with irradiated infective L3 reduces the worm load. This present study investigated whether the additional activation of cytosolic nucleic acid receptors as adjuvant improves the efficacy of a vaccination with irradiated L3 larvae of the rodent filaria *Litomosoides sigmodontis*. Subcutaneous injection of irradiated L3 larvae in combination with poly(I:C) or 3pRNA resulted in increased neutrophil recruitment to the skin, accompanied by higher IP-10/CXCL10 and IFN-β RNA levels at the site of injection. To investigate the *in vivo* impact on parasite clearance, BALB/c mice received 3 subcutaneous injections in 2-week intervals with irradiated L3 larvae in combination with poly(I:C) or 3pRNA prior to the challenge infection. Serum analysis before the challenge infection confirmed the induction of *L. sigmodontis*-specific antibodies in response to the immunization and serum from immunized mice significantly reduced larval motility *in vitro* with naïve cells. 63 days after the challenge infection, vaccination with irradiated L3 larvae in combination with poly(I:C) or 3pRNA led to a significantly greater reduction in adult worm counts by 73% and 57%, respectively, compared to the immunization with irradiated L3 larvae alone (45%). Further, the treatment of *L. sigmodontis* infection with 3pRNA alone, but not poly(I:C), resulted in a reduced worm burden, supporting the therapeutic potential for the activation of RIG-I with 3pRNA. In conclusion, our data demonstrate that the additional activation of nucleic acid sensing immune receptors boosts the immune response and provides better protection against *L. sigmodontis*. Thus, the use of nucleic acid receptor agonists as vaccine adjuvants represents a promising novel strategy to improve the efficacy of vaccines against filariae and potentially of other helminths.

**Author Summary:** Filarial nematodes can cause debilitating diseases such as onchocerciasis and lymphatic filariasis that present a major public health burden in the tropics and subtropics, putting more than a billion people at risk of infection. Filarial diseases are transmitted to humans by insect vectors as they take a blood meal. The WHO classifies both filarial infections as neglected tropical diseases and aims to eliminate the transmission of onchocerciasis and eliminate lymphatic filariasis as public health problem by 2030. However, up to date there is no vaccination available that could support the efforts to eliminate filarial diseases and potentially helminth infections in general. Here, we used the well-established murine model for filarial infection, *Litomosoides sigmodontis*, to test the use of nucleic acid receptor agonists as vaccine adjuvants to enhance local immune responses. We found that infection with *L. sigmodontis* induces type I IFN and our vaccine strategy enhances the production of type I IFN resulting in increased parasite-specific immune responses and enhanced worm clearance. In summary, our study provides a promising novel approach for a vaccination strategy using cytosolic RNA receptor agonists.

## Introduction

Helminth infections affect more than 1 billion people and put a high socio-economic burden on endemic countries (1). Filariae are extraintestinal helminths, which can cause debilitating diseases such as lymphatic filariasis (elephantiasis) and onchocerciasis (river blindness). Currently there are 859 million (2) and 218 million (3) people living in areas of ongoing transmission of these diseases, respectively. The elimination of both diseases until 2030, at least in a considerable percentage of countries, was stated as goal of the World Health Organization (WHO) (4, 5). The control of both diseases is mainly mediated by annual mass drug administrations (MDA) of drugs that temporarily inhibit filarial embryogenesis and the transmission of the disease, but do not kill the adult filariae (6, 7). To support elimination, the development of a potent prophylactic and therapeutic vaccine would be a powerful tool. However, currently there is no vaccination available for any human helminth infection (8, 9) and helminth vaccine candidates are providing only limited protection. For filarial infections, one of the best efficacies is obtained when irradiated L3 larvae are used for vaccinations, which can be used to study protective immune responses and explore novel vaccination approaches.

In this project we used the *Litomosoides sigmodontis* murine model to identify new vaccination strategies. *L. sigmodontis* establishes chronic infections in susceptible BALB/c mice and triggers immune responses that resemble those of human filarial infections (10). The parasite is transmitted to the vertebrate host via the bite of the tropical rat mite *Ornithonyssus bacoti*. After transmission, the infective L3 larvae migrate from the infection site in the skin via the lymphatic system and the pulmonary blood circulation to the pleural cavity where they molt into L4 larvae around day eight and further into adult worms by day 30 post infection. Starting around 50 days post infection the adult female worms release their offspring, microfilariae (MF), into the peripheral blood (11). Previous studies showed that a triple immunization with irradiated *L. sigmodontis* L3 larvae every week resulted in the generation of parasite-specific IgG1 antibodies and a reduction in adult worm burden following a challenge infection of around 50% (12–15).

One of the challenges to develop vaccines against helminth infections is the modulation of the host’s immune system by the helminths. During filarial infections, the host’s immune system is strongly regulated by the parasite in order to facilitate its survival in the host (16, 17). Type 2-associated immune responses have been linked to protection (18), however also a type 1/type 2 balanced immune response appears to be crucial for parasite elimination (9, 19–21). To overcome those protective immune responses, helminths release immunomodulatory molecules and establish a regulatory, anti-inflammatory immune milieu over time (16–18, 22–24). So far, a possible contribution of nucleic acid immunity in the context of helminth infection has not been addressed. Nucleic acid immunity is best studied in the context of viral infections where it triggers dominant type 1-associated immune responses (25). Different RNA- (e.g. RIG-I, MDA5; TLR7/8) and DNA- (e.g. via STING; TLR9) sensors in the cytosol or endosome of eukaryotic cells (26) activate downstream cascades that involve the induction of a type I interferon (IFN) response (26) which provides innate anti-viral immunity. A possible impact of type I IFN responses on filarial infections is unknown.

In this study we report the induction of type I IFN *in vivo* during the infection with *L. sigmodontis*. Further, we observed a significant reduction in worm burden after the treatment with the specific RIG-I ligand 3pRNA. In addition, we show that the use of nucleic acid receptor agonists as vaccine adjuvant enhances local immune responses including the production of type I IFN during immunization, which results in increased parasite-specific immune responses and enhanced worm clearance after infectious challenge. Consequently, we propose a protective role for type I IFN signaling during filarial infections and present the use of the cytosolic RNA receptors agonists as novel strategy to improve filarial vaccination efficacy.

## Results

### L. sigmodontis infection triggers IFN-β production in vivo

In order to study a potential role of type I IFNs during filarial infections, albino C57BL/6 IFN-β Luciferase reporter mice were infected with *L. sigmodontis*. The infection induced significantly increased IFN-β transcripts by pleural cells at 5, 12 and 30 days after the infection (Figure 1A), which correlated with the arrival of L3s in the thoracic cavity, the molting into L4 stage larvae and adult worms, respectively. During the course of infection, transcription of IFN-β increased (Figure 1A). The activation of nucleic acid receptors by *in vitro* stimulation of pleural cells with poly(I:C) further increased the IFN-β production (Figure 1A). While at 5 days after the infection the IFN-β production showed no correlation with the *L. sigmodontis* worm burden (Figure 1B), a significant positive correlation between the worm burden and IFN-β production was observed at 12 and 30 days after the infection (Figure 1C+D).

**Fig 1:**
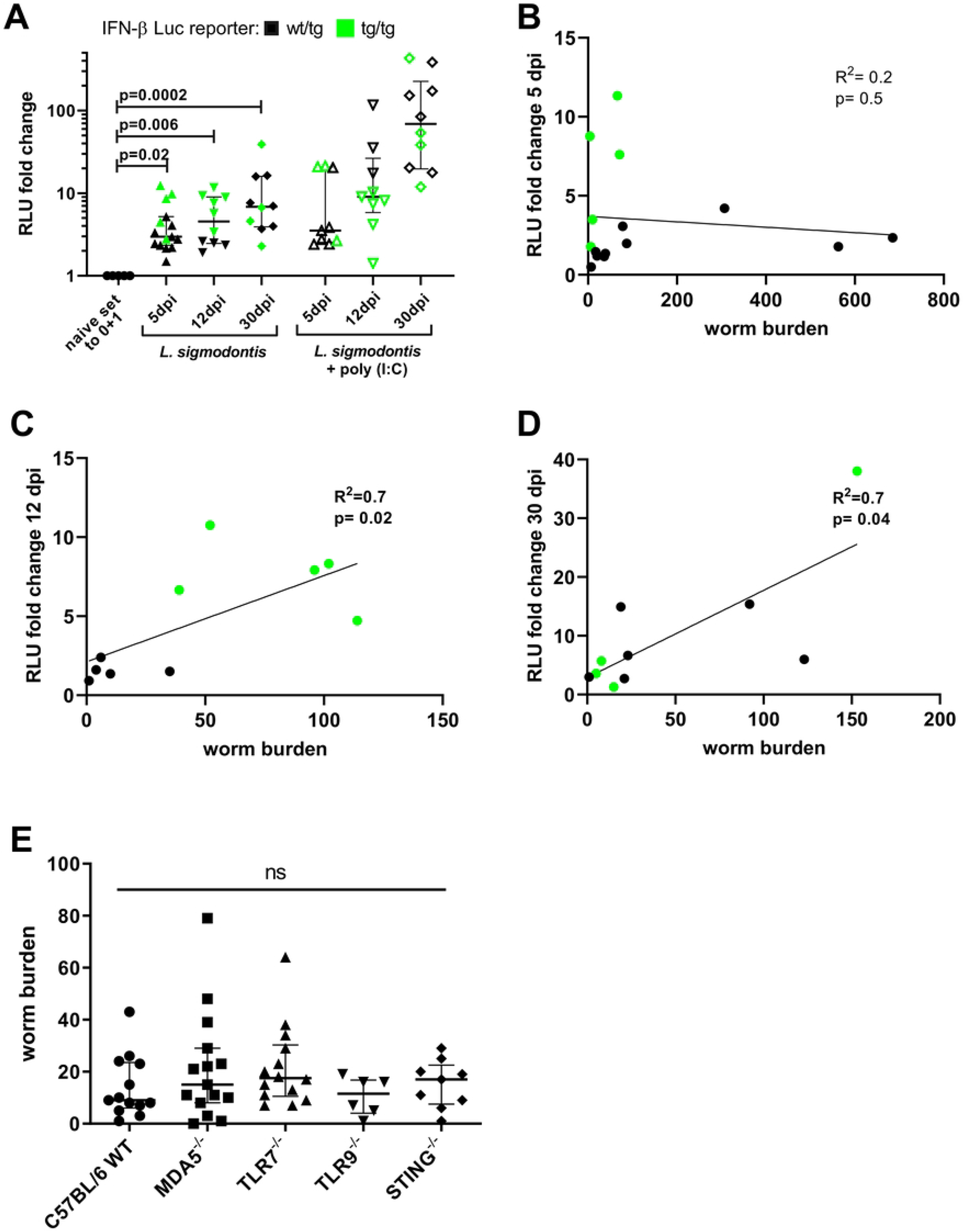
*In vivo* IFN-β production in pleura cells positively correlates with *L. sigmodontis* infection duration and worm burden. **(A-D)** Pleural cells were isolated from IFN-β luciferase reporter mice at 5, 12 and 30 days after the infection with *L. sigmodontis*. IFN-β transcripts as fold change was calculated based on the average relative light unit (RLU) measured in naïve control animals. The naïve values were redefined as 0 +1 to allow log scale **(A)** IFN-β transcripts by pleural cells at 5, 12 and 30 days post infection (dpi) with or without additional poly(I:C) stimulation. **(B-D)** Correlation of IFN-β and worm burden per animal at **(B)** 5 dpi, **(C)** 12 dpi and **(D)** 30 dpi. **(E)** *L. sigmodontis* worm burden of C57BL/6 wild-type (WT), MDA5^-/-^, TLR7^-/-^, TLR9^-/-^, and STING^-/-^ mice at 14 dpi. **(A+E)** Error bars show median with IQR. Data were statistically analyzed using Kruskal-Wallis with Dunn’s post-hoc test. **(B-D)** Spearman correlation including linear regression line is shown. **(A-D)** Data from 5 dpi are pooled from three individual experiments and data from 12 dpi and 30 dpi are pooled from two individual experiments with n=9-15. **(E)** Data for C57BL/6 WT, MDA5^-/-^ and TLR7^-/-^ were pooled from 2 individual experiments with n=6-15. **(A-E)** Data were pooled after failing Spearman’s rank correlation test for heteroscedasticity.

### Unaltered larval migration in nucleic acid receptor deficient mice

Given the increasing production of IFN-β during *L. sigmodontis* infection, the impact of nucleic acid receptors on larval migration was assessed upon natural *L. sigmodontis* infection of C57BL/6 wildtype (WT), MDA5^-/-^, TLR7^-/-^, TLR9^-/-^ and STING^-/-^ mice. Since larval migration is completed by day eight (27) and C57BL/6 start clearing *L. sigmodontis* infections around day 35 (28–30), the worm burden was assessed twelve days after infection. No changes in the pleural worm burden were determined for any of the knockout strains analyzed (Figure 1E). Lack of TLR3 has been previously described to have no influence on *L. sigmodontis* worm burden (31). These findings indicate that *L. sigmodontis* infection induces IFN-β production, but the lack of single nucleic acid receptors does not lead to an enhanced or reduced susceptibility for filarial infection.

### Co-immunization with poly(I:C) or 3pRNA enhances local immune responses to irradiated L3 larvae

Based on our findings that *L. sigmodontis* infection induces type I IFN production and reports showing that filariae modulate nucleic acid sensing pathways (32–35), we hypothesized that the activation of nucleic acid receptors might enhance worm clearance and the use of agonists as vaccine adjuvants might increase vaccination efficacy. To that end the immunostimulatory potential of various nucleic acid receptor agonists was analyzed four hours after subcutaneous injection (S1 Tab.). The injection of all agonists but R848 resulted in increased frequencies of CD11b^+^ cells in the skin. Only the injection of poly(I:C) or 3pRNA, but not R848 or CpG-C, induced a strong local type I IFN response. Therefore, poly(I:C) and 3pRNA were selected for further analysis. Poly(I:C) is a known agonist of the RNA sensors TLR3, MDA5 and RIG-I (36, 37), while 3pRNA activates RIG-I (38).

Poly(I:C) and 3pRNA were administered as adjuvant for the immunization with att. *L. sigmodontis* L3 larvae and local as well as systemic immune responses were analyzed four hours after the immunization (Figure 2A). Flow cytometric analysis revealed a significantly increased frequency of neutrophils in the skin after the injection of att. L3 larvae with poly(I:C) or 3pRNA, compared to the 0.9% NaCl control or injection of att. L3 larvae alone (Figure 2B). At the same time, there was a significantly decreased frequency of monocytes and CD11b^+^ DCs in the skin of mice injected with att. L3 larvae and poly(I:C) or 3pRNA, compared to the 0.9% NaCl control or injection of att. L3 larvae alone (Figure 2C + D). Levels of IFN-β and IP-10 (CXCL10) within the skin were increased in response to the injection of att. L3 larvae with poly(I:C) or 3pRNA, compared to the 0.9% NaCl control (Figure 2E + F). The injection of att. L3 alone did not induce local IFN-β or IP-10 (CXCL10) responses. None of the injections resulted in increased systemic responses of the chemokine IP-10 (CXCL10) (Figure 2G). Taken together, poly(I:C) and 3pRNA enhanced local immune responses when co-administered with att. L3 larvae for immunization.

**Fig 2:**
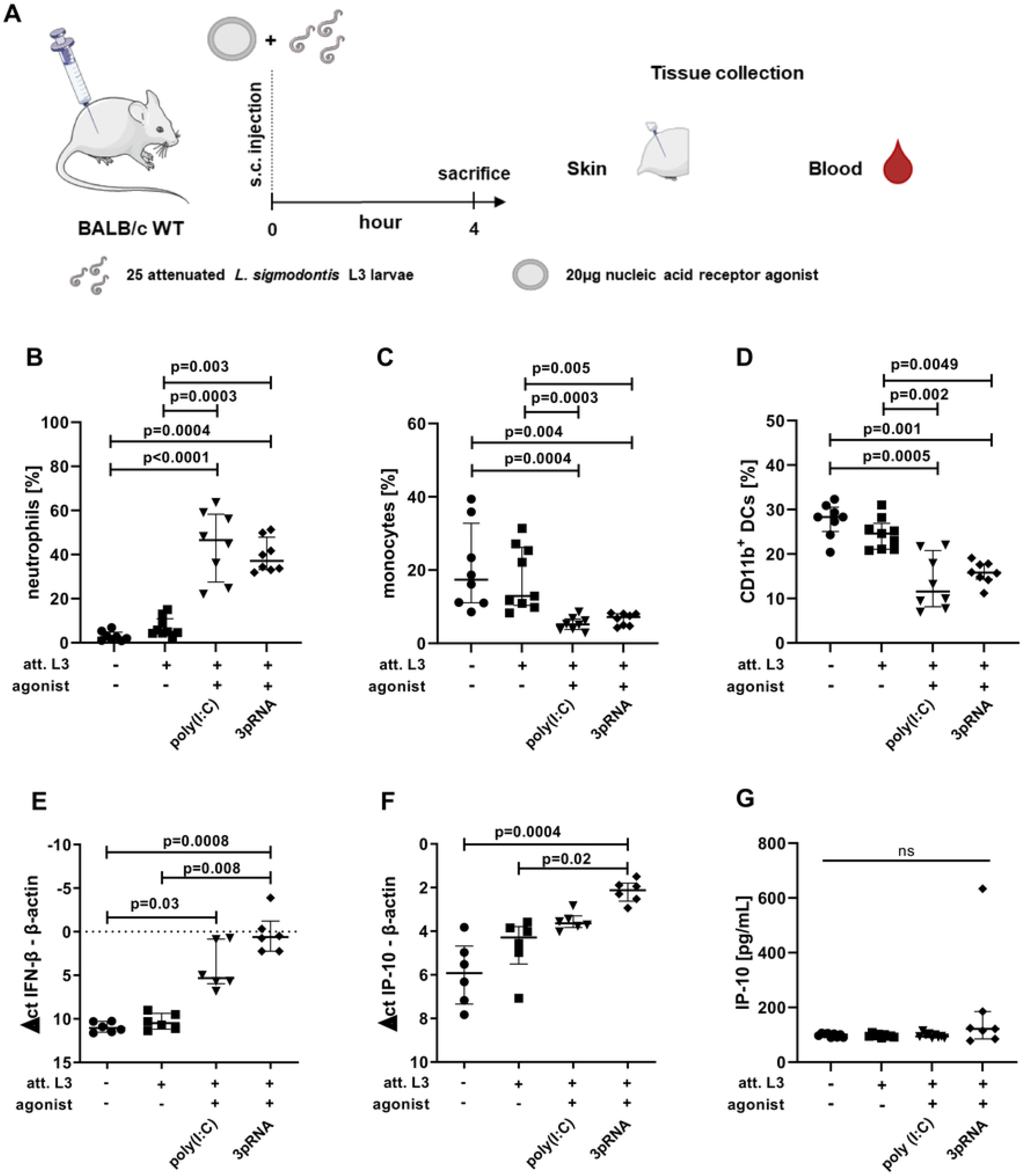
Enhanced local immune activation after injection of poly(I:C) or 3pRNA. **(A-G)** Mice were injected subcutaneously with attenuated (att.) *L. sigmodontis* L3 larvae with or without 3pRNA and poly(I:C)) and the skin was analyzed four hours after injection. **(A)** Experimental setup. **(B-D)** Skin cells were analyzed by flow cytometry. Frequency of **(B)** neutrophils (CD45^+^CD11b^+^Ly6G^+^), **(C)** monocytes (CD45^+^CD11b^+^Ly6C^+^Ly6G^-^), **(D)** CD11b^+^ DCs (CD45^+^CD11b^+^Ly6C^-^Ly6G^-^CD11c^+^SiglecF^-^) among CD45^+^ cells. **(E-F)** Skin samples were analyzed by RT-PCR. Δct values of **(E)** IFN-β expression and **(F)** IP-10/CXCL10 expression compared to β-actin levels in the corresponding sample. **(G)** Serum concentrations of IP-10/CXCL10 quantified by ELISA. **(B-G)** Error bars show median with IQR. Data were statistically analyzed by Kruskal-Wallis with Dunn’s post-hoc test. **(B-D)** Data from one experiment, n=8. **(E+F)** representative for three experiments with n=5-6. **(G)** Data pooled from two individual experiments with n=7-9.

### 3pRNA treatment reduces larval burden

In order to clarify whether the enhanced immune activation in response to poly(I:C) and 3pRNA improves protection during *L. sigmodontis* infection, mice were subcutaneously injected with poly(I:C) or 3pRNA prior to s.c. infection with 25 *L. sigmodontis* L3 larvae, followed by an intravenous injection of poly(I:C) or 3pRNA (Figure 3A). Nine days after infection the larval count in the pleural cavity was significantly reduced in animals treated with 3pRNA (p=0.03) compared to control animals (Figure 3B). After the treatment with poly(I:C) or 3pRNA the total cell count in the pleura cavity was reduced compared to control animals (Figure 3C). The neutrophil count remained similar in all groups (Figure 3D). However, neutrophils in the pleura of mice treated with either poly(I:C) or 3pRNA were significantly more activated, based on the expression of I-Ab and CD86 (Figure 3E + F). Eosinophil counts were lower in mice treated with poly(I:C) or 3pRNA (p=0.04) compared to control animals (Figure 3G). This was accompanied by a significant reduction in M2-like macrophage numbers after the treatment with poly(I:C) (p=0.007) or 3pRNA (p=0.04) compared to control animals, assessed by intracellular RELMα expression (Figure 3H). At the same time the counts of M1-like macrophages were increased in the pleura of animals treated with poly(I:C) or 3pRNA (p=0.04), when compared to control animals (Figure 3I). Also, the pleural cavity counts of CD11b^+^ DCs, B cells and CD4^+^ T cells were lower in animals treated with poly(I:C) or 3pRNA, while CD8^+^ T cell and NK cell counts remained similar to control animals (S1 Fig). Taken together, the treatment with nucleic acid receptor agonist 3pRNA alone improves *L. sigmodontis* clearance *in vivo* while treatment with poly (I:C) and 3pRNA alone lead to changes in the immune cell composition within the pleural cavity.

**Fig 3:**
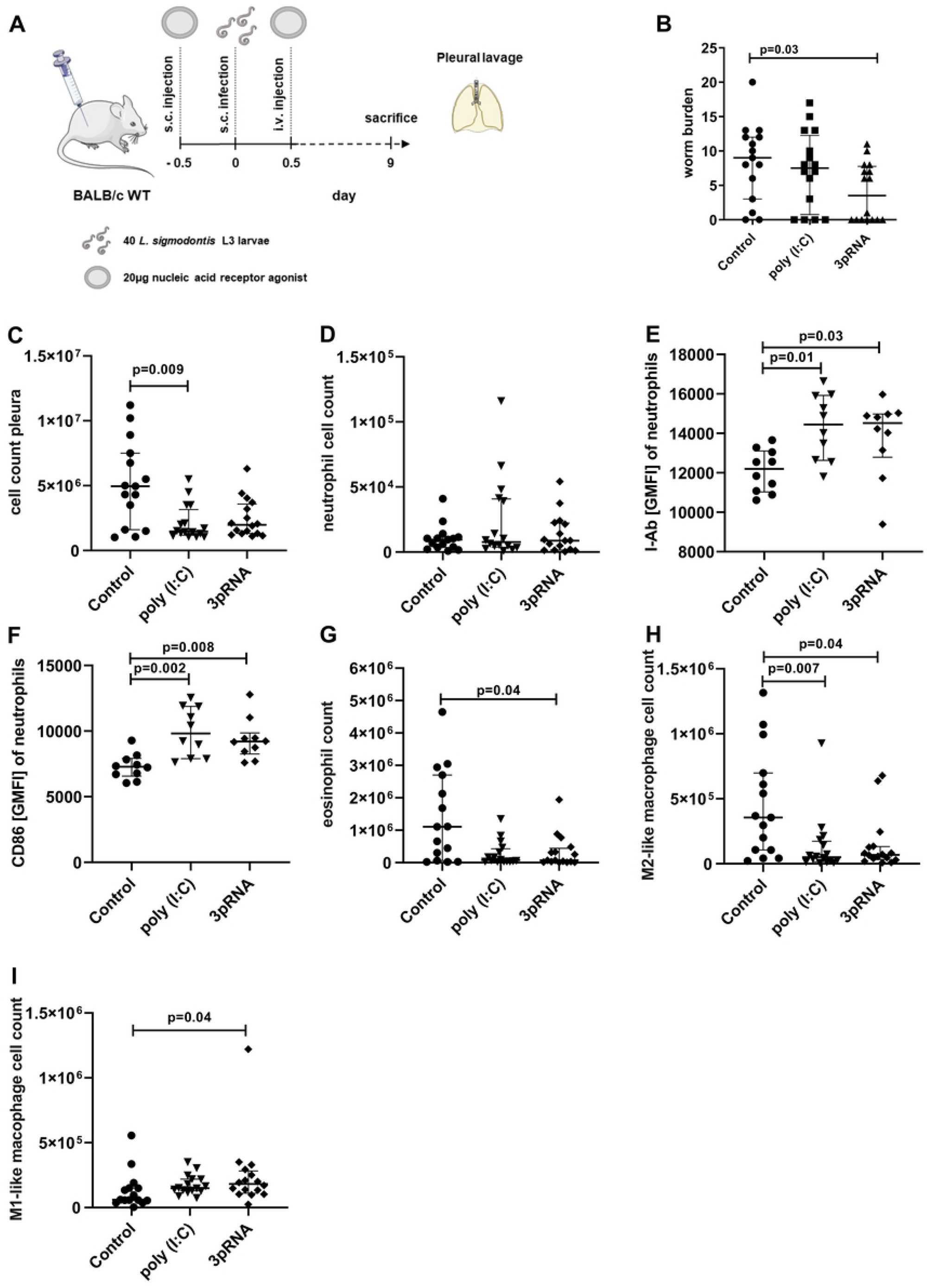
Reduced worm burden after treatment with poly(I:C) or 3pRNA. **(A-I)** Mice were subcutaneously (s.c.) injected with poly(I:C) or 3pRNA. 12 hours later, mice were s.c. infected with 40 *L. sigmodontis* L3 larvae and intravenously injected with the same agonist 12 hours after the infection. Analysis was performed nine days after infection. **(A)** Experimental setup. Quantification of **(B)** *L. sigmodontis* larvae and **(C)** the total cell count in the pleural cavity. **(D-I)** Pleural cells were analyzed by flow cytometry. **(D)** Number of pleural neutrophils (SiglecF^-^ CD11b^+^CD11c^-^Ly6C^int^ Ly6G^+^) and their expression of **(E)** I-Ab and **(F)** CD86. Number of pleural **(G)** eosinophils (SiglecF^+^), **(H)** M2-like macrophages (SiglecF^-^CD11b^+^CD11c^-^Ly6G^-^ Ly6C^int^RELMα^+^) and **(I)** M1-like macrophages (SiglecF^-^CD11b^+^CD11c^-^Ly6G^-^Ly6C^int^RELMα^-^). **(A-I)** Data shown as median with IQR. Data were statistically analyzed by Kruskal-Wallis with Dunn’s post-hoc test. **(A+B)** Data pooled from two individual experiments. **(D-I)** Data from one experiment, representative for two individual experiments with n=15-16.

### Immunization with adjuvants enhances functional parasite-specific antibody responses

Since the treatment with poly(I:C) or 3pRNA enhanced filarial clearance, the agonists were included to the *L. sigmodontis* immunization strategy. To that end mice were subcutaneously immunized with att. L3 larvae in combination with poly(I:C) or 3pRNA every two weeks for a total of three times (Figure 4A). This was followed by a natural challenge infection two weeks after the last immunization. The serum obtained two weeks after full immunization, but prior to challenge infection, was analyzed for parasite-specific antibodies. Immunization with att. L3 larvae alone or along with poly(I:C) or 3pRNA induced the production of *L. sigmodontis*-specific IgE compared to non-immunized mice (Figure 4B). All immunization regimes resulted in significant production of *L. sigmodontis*-specific IgG1 (Figure 4C). The immunization with att. L3 larvae alone did not induce significant levels of *L. sigmodontis*-specific IgG2a/b (Figure 4D). However, there was a significant IgG2a/b response in animals immunized with a combination of att. L3 with non-formulated poly(I:C), poly(I:C) or 3pRNA. The use of non-formulated poly(I:C) or poly(I:C) as adjuvant also significantly increased the IgG2a/b production compared to the immunization with att. L3 larvae alone.

**Fig 4:**
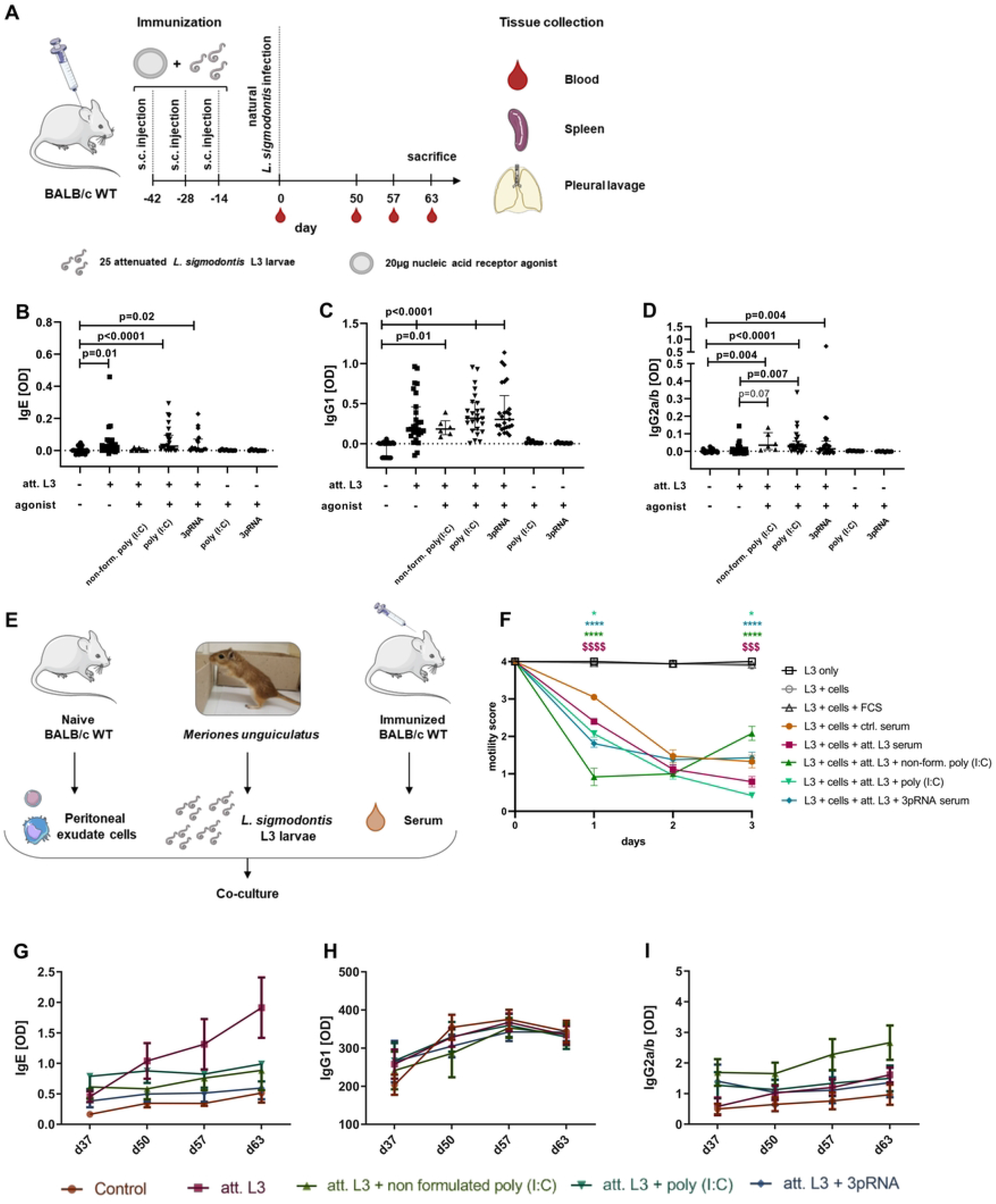
Immunization with adjuvants enhances functional parasite-specific antibody responses. **(A-D)** Mice were immunized for three times in two-week intervals by subcutaneous injection of attenuated (att.) *L. sigmodontis* L3 larvae in combination with non-formulated poly(I:C) (p(I:C)), p(I:C) or 3pRNA. **(A)** Experimental setup. **(B-D)** *L. sigmodontis*-specific **(B)** IgE, **(C)** IgG1 and **(D)** IgG2a/b serum antibody levels two weeks after the last injection and before challenge infection. **(E)** Experimental setup of an ADCC assay that was performed using co-cultures with serum of immunized BALB/c mice as well as *L. sigmodontis* L3 larvae and naïve peritoneal exudate cells. The motility of individual larvae shown in **(F)** was assessed for three days by the following score: 4: fast and continuous movement, 3: slower but continuous movement, 2: slower and discontinuous movement, 1: sporadic movement limited to the ends, 0: no movement. **(G-H)** Immunized mice were naturally infected with *L. sigmodontis* two weeks after the last immunization injection. Serum was collected 37, 50, 57 and 63 days after infection and analyzed by ELISA for *L. sigmodontis*-specific **(G)** IgE, **(H)** IgG1 and **(I)** IgG2a/b antibodies. **(B-D)** Data shown as median with IQR. Statistical analysis using Kruskal-Wallis with Dunn’s post-hoc test with n=8-28. Data from untreated mice and groups receiving att. L3 larvae with or without agonist were pooled from 3 individual experiments. Data from att. L3 with non-formulated poly(I:C) and groups that only received agonist are from one experiment. **(F)** Data presented as mean ± SEM and was statistically analyzed by a 2-way ANOVA with Bonferroni’s multiple comparison test. $$$$ p<0.0001 comparison of L3 + att. L3 serum to L3 + ctrl. serum. * p<0.05, *** p<0.0001: comparison of groups including agonist treatment to L3 + att. L3 serum. Data from L3 + cells + FCS pooled from two independent experiments. Data from L3 + att. L3 + non-formulated poly(I:C) serum from one experiment. Other data pooled from three individual experiments. For L3 + att. L3 + non-formulated poly(I:C) serum with n=12 larvae. For other groups n=49 - 72 larvae. **(G-I)** Data from one experiment presented as mean ± SEM with n=6-10.

In order to assess functionality, an antibody-dependent cellular cytotoxicity assay was performed. Since the immune cell counts of the naïve pleural cavity is too low to perform ex vivo assays, naïve peritoneal cells, mainly consisting of myeloid and B cells (S2 Fig), were co-cultured with *L. sigmodontis* L3s, the medium was supplemented with the serum of immunized mice and the motility of L3s was scored over time (Figure 4E). The supplementation with serum of unimmunized mice led to a continuous slow motility of L3s (score 3) after one day, and discontinuous movements in part restricted to the ends of the L3s at two and three days of culture (score 1.3, Figure 4F). The supplementation with serum from mice immunized with att. L3 larvae alone resulted in significantly reduced larvae motilities on day one (score 2.4) and three (score 0.8) of culture. Compared to this, the serum of animals immunized with a combination of att. L3 larvae and non-formulated poly(I:C), poly(I:C) or 3pRNA resulted in significantly reduced larval motility on day one (score 0.9 for non-formulated poly(I:C), score 2 for poly(I:C), score 1.8 for 3pRNA). On day three only the group with att. L3 and poly(I:C) immunized serum had a motility score that was significantly lower (score 0.4) compared to the effect of serum from mice immunized with att. L3 alone. In co-cultures with serum from animals immunized with att. L3 larvae and non-formulated poly(I:C) the larval motility recovered to a score of 2.1.

Parasite-specific antibodies were further measured on 37, 50, 57, and 63 days after challenge infection. At all times parasite-specific IgE was lowest in the unimmunized control group (Figure 4G). In all groups the IgE level remained similar during the infection, with the exception of mice immunized with att. L3 larvae alone, which showed increased IgE values on day 63 after the infection. IgG1 levels were lowest in unimmunized mice on day 37 after infection, but highest at the following time points (Figure 4H). IgG1 levels were similar in all immunized groups and remained stable during the entire time course. IgG2a/b levels were lowest in non-immunized animals (Figure 4I). The immunization with att. L3 larvae alone led to increasing IgG2a/b levels during the course of infection, whereas the combination immunization of att. L3 larvae with non-formulated or poly(I:C) resulted in higher IgG2a/b levels at days 37, 50 and 57 after challenge infection. Overall, the immunization with att. L3 larvae induced parasite-specific, functional antibodies, which was enhanced by the use of nucleic acid receptor agonists as adjuvant.

### Immunization with poly(I:C) or 3pRNA as adjuvant significantly reduces the worm burden following challenge infection in mice

63 days after the challenge infection of immunized mice the adult worm burden was quantified (Figure 5A). The immunization with att. L3 larvae alone resulted in a worm burden reduction of 45% compared to unimmunized mice. The implementation of non-formulated poly(I:C) as adjuvant did not improve worm clearance (39% worm burden reduction). Poly(I:C) as immunization adjuvant resulted in a significant reduction in worm burden of 73% when compared to unimmunized mice (p=0.0002) and by trend when compared to the immunization with att. L3 larvae alone (p=0.06). The use of 3pRNA in combination with att. L3 larvae for immunization resulted in a significant reduction in worm burden by 57% compared to unimmunized animals (p=0.0008). No changes were observed regarding the sex of the worms. (S3 Fig). Of note, none of the immunization regimens led to a complete absence of MF-positive mice (Table 1), indicating that remaining filariae are viable, fertile and potentially able to maintain filarial transmission. However, the percentage of MF-positive animals in the control group was 50% and this was reduced to 33.3% by the immunization with att. L3 larvae alone, and further reduced to 29.4% and 16.7% by the use of poly(I:C) or 3pRNA as agonist, respectively.

**Table 1.** Immunization including poly(I:C) or 3pRNA adjuvant reduces the percentage of animals developing microfilaremia. Mice were immunized three times in two-week intervals by subcutaneous injection of attenuated (att.) *L. sigmodontis* L3 larvae in combination with non-formulated poly(I:C) (p(I:C)), p(I:C) or 3pRNA. Two weeks after the last injection, the mice were naturally infected with *L. sigmodontis*. 63 days after the infection microfilariae (MF) were quantified in the peripheral blood. The percentage of MF-positive animals per group was determined, n=6-20. Pooled data from two individual experiments, data on non-formulated p(I:C) from one experiment.

| Group | animals<br>total [#] | MF<sup>+</sup> animals<br>[#] | MF<sup>+</sup> animals<br>[%] |
| --- | --- | --- | --- |
| Control | 20 | 10 | 50% |
| att. L3 | 18 | 6 | 33.3% |
| att. L3 + non-formulated poly(I:C) | 6 | 3 | 50% |
| att. L3 + poly (I:C) | 17 | 5 | 29.4% |
| att. L3 + 3pRNA | 18 | 3 | 16.7% |

**Fig. 5:**
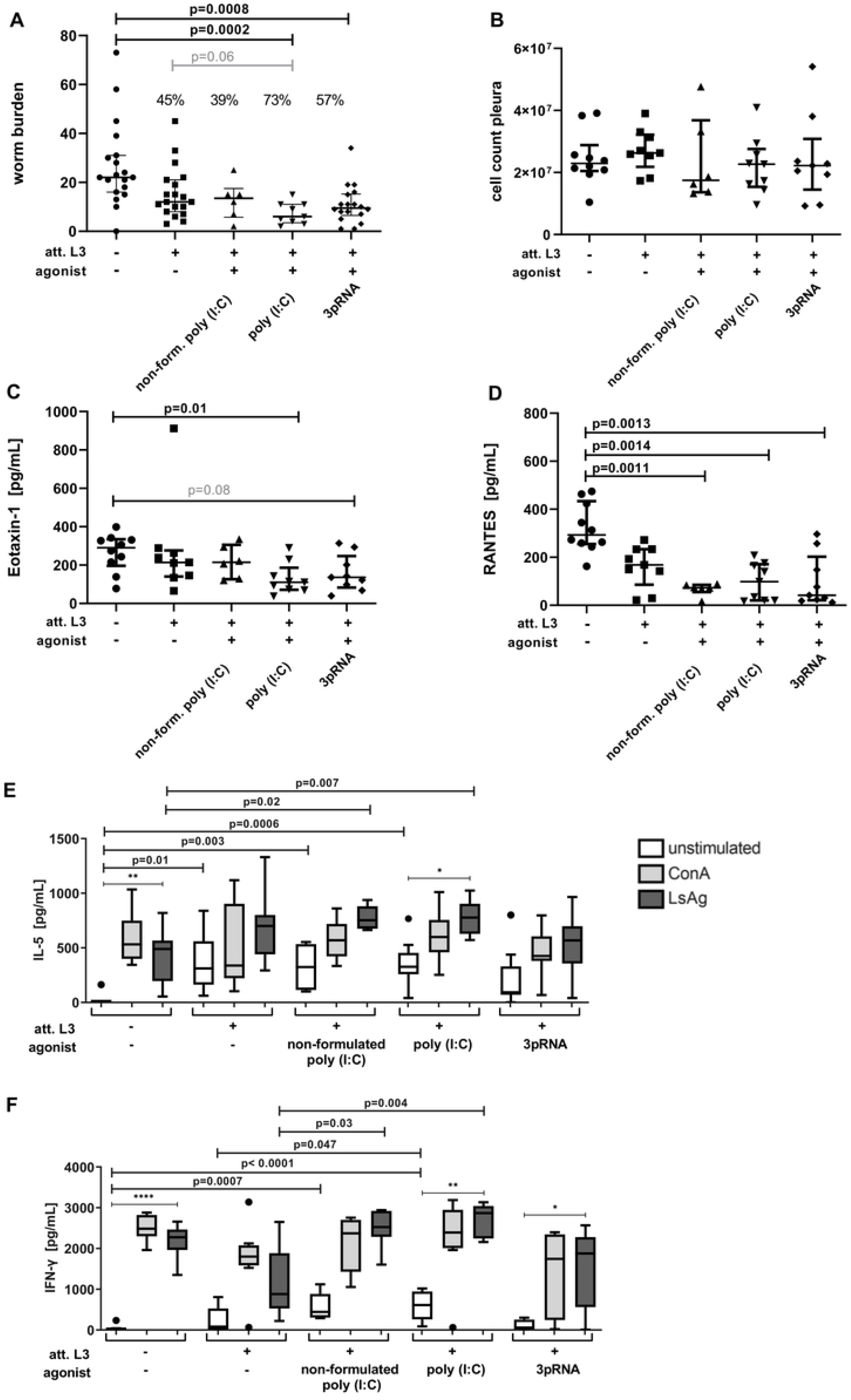
Immunization with poly(I:C) or 3pRNA as adjuvant significantly reduces the worm burden following challenge infection in mice. **(A-F)** Mice were immunized three times in two-week intervals by subcutaneous injection of attenuated (att.) *L. sigmodontis* L3 larvae in combination with non-formulated poly(I:C) (p(I:C)), p(I:C) or 3pRNA. Two weeks after the last injection, the mice were naturally infected with *L. sigmodontis* for 63 days. **(A)** Quantification of adult worms and **(B)** total cell count in the pleural cavity. Quantification of **(C)** Eotaxin-1/CCL11 and **(D)** RANTES/CCL5 in the pleura lavage by ELISA. **(E)** IL-5 and **(F)** IFN-γ concentrations of splenocytes that were restimulated with Concanavalin A (ConA) or *L. sigmodontis* adult worm extract (LsAg). **(A-D)** Data shown as median with IQR. **(E+F)** Data shown as box and whiskers blot. **(A-F)** Data were statistically analyzed by Kruskal-Wallis with Dunn’s post-hoc test. **(A)** Pooled data from two individual experiments; data on non-formulated poly(I:C) experiment from one experiment. **(B)** Data from one experiment **(C-F)** and representative for two individual experiments except data on non-formulated poly(I:C) with n=6-20.

Despite the reduction in worm burden there were no notable changes in the pleural cell count in any immunized group compared to the control animals (Figure 5B). The concentration of the chemokine CCL11/eotaxin-1 was significantly reduced (p=0.01) in the pleural cavity of mice immunized with a combination of att. L3 larvae and poly(I:C) and reduced in the pleural cavity of mice immunized with att. L3 larvae and 3pRNA (Figure 5C; p=0.08). The concentration of the chemokine CCL5 (RANTES) was significantly decreased in the pleural cavities of mice immunized with a combination immunization of att. L3 larvae with non-formulated poly(I:C), poly(I:C) or 3pRNA, compared to unimmunized mice (Figure 5D).

Given that local immune responses in the pleural cavity may be affected by the worm burden, systemic effects, i.e. the cytokine response of spleen cells were additionally quantified. Unstimulated splenocytes from mice immunized with att. L3 alone or a combination with non-formulated poly(I:C) or poly(I:C) released significantly higher levels of IL-5 than cells from unimmunized mice (Figure 5E). The IL-5 response to the restimulation with LsAg was significantly enhanced in spleen cells from mice immunized with non-formulated poly(I:C) or poly(I:C), in comparison to spleen cells from unimmunized mice. At the same time we found a significantly increased IFN-γ release by splenocytes from animals immunized with a combination of att. L3 larvae and non-formulated poly(I:C) or poly(I:C) compared to cells from unimmunized mice (Figure 5F). Further, significantly higher IFN-γ responses to the restimulation with LsAg in mice immunized with att. L3 larvae and non-formulated poly(I:C) or poly(I:C), compared to the immunization with att. L3 larvae alone were observed.

Taken together, the use of the nucleic acid receptor agonists poly(I:C) or 3pRNA significantly reduced the adult worm burden, increased parasite-specific immune responses and enhanced the immunization efficacy.

## Discussion

In this study we demonstrate that type I interferon is increasingly produced during infection with *L. sigmodontis* at the site of the parasite. The subcutaneous injection of att. L3 larvae along with nucleic acid receptor agonists poly(I:C) and 3pRNA enhanced local immune responses. Importantly, the treatment of mice with 3pRNA alone at the time of *L. sigmodontis* infection resulted in significantly reduced worm burden implicating an innate anti-helminth effect following RIG-I activation. Furthermore, the implementation of poly(I:C) and 3pRNA as adjuvants during the immunization with irradiated *L. sigmodontis* L3 larvae was successful in enhancing parasite-directed immune responses during subsequent infection, resulting in a reduced adult worm burden of 73% and 57%, respectively.

These results demonstrate that nucleic acid receptor agonists such as poly(I:C) and 3pRNA improve the efficacy of a vaccine against the filarial nematode *L. sigmodontis* and may present a novel strategy to booster the efficacy of helminth vaccines that are currently developed. Our data revealed that infection with *L. sigmodontis* induced the production of type I IFNs at the site of parasite manifestation. Type I IFN production is a downstream event after the activation of nucleic acids receptors and is in general associated with a Th1-dominated immune response (26). However, in the context of *Schistosoma mansoni* infection, type I IFNs were crucial to drive Th2 cell induction (39) and may indicate that a balanced Th1/Th2 immune response is necessary for protection and that type I IFNs may support such protective immune responses. Other studies showed that helminths modulate nucleic acids sensing (32–35), for example by the downregulation of TLR3 and TLR7 expression by *B. malayi* MF (32, 33). Further, it was shown that *B. malayi* L3 larvae initially elicit type 1-associated responses, but during the course of infection actively suppress these (40, 41). This might explain why there was no impact on larval migration during the early time point of *L. sigmodontis* infection in the tested nucleic acid receptor-deficient mouse strains. Alternatively, redundant protective immune responses may be triggered by *L. sigmodontis* and deficiency of a single nucleic acid receptor may not be sufficient to overcome these protective mechanisms. Of note, the nucleic acid receptor deficient mouse strains as well as the IFN-β Luciferase reporter mice were on a C57BL/6 background, which is semi-susceptible for the infection with *L. sigmodontis*, as the filariae are cleared in the C57BL/6 mouse strain shortly after the molt into the adult stage (19). Thus, it would be of interest to analyze the type I IFN response during the course of infection in BALB/c mice, which are susceptible for the *L. sigmodontis* infection and allow the development of chronic infections with MF detectable in the peripheral blood (10).

This BALB/c mouse strain was used for the subsequent experiments analyzing the potential of nucleic acid receptor ligands as inducers of protective immune responses. The agonist poly(I:C) mainly targets the cytosolic RNA sensors MDA5 and RIG-I, but is also known to activate the endosomal RNA sensor TLR3 (36, 37), while the agonist 3pRNA specifically targets RIG-I (38). Upon subcutaneous injection of the nucleic acid receptor agonists poly(I:C) or 3pRNA alone there was a rapid influx of neutrophils. Neutrophils are major effector cells that support the elimination of invading L3 larvae (42, 43). Therefore, it is likely that due to the treatment with poly(I:C) or 3pRNA part of the L3 larvae were already eliminated in the subcutaneous tissue. Further, it has been shown that type I IFNs enhance the migration of DCs to the draining lymph node and improve their co-stimulatory potential (44). In line with this, local type I IFN responses were triggered upon *L. sigmodontis* infection and DC frequencies decreased after agonist injection. Therefore, the local activation with poly(I:C) and 3pRNA might enhance the activation of innate but also adaptive immune responses targeting the invading L3 larvae. The treatment with 3pRNA resulted in a significant reduction of L3 larvae reaching the pleural cavity, demonstrating that the activation of the RIG-I pathway by itself can induce protective innate immunity. Treatment with poly(I:C) did not show such a reduction in larval recovery.

In the context of a vaccination with irradiated L3 larvae, poly(I:C) as well as 3pRNA enhanced the efficacy of the immunization. Previous studies showed that the immunization with irradiated *L. sigmodontis* L3 larvae results in the generation of parasite-specific antibodies (13), a reduction in adult worm burden of around 50% (12, 14, 45, 46) and a reduction of MF positive animals (13). In contrast to previous studies that used subcutaneous injections of L3 larvae for the challenge infection, we used a natural infection via the mite vector. Despite this difference, our study observed a reduction in adult worm burden of 45% for mice that solely received the immunization with irradiated L3 larvae. Furthermore, the induction of parasite-specific IgG1 and IgG2 antibodies and the reduction in MF-positive animals was replicated in our study. The addition of poly(I:C) or 3pRNA to the vaccination regimen with irradiated L3 larvae enhanced the production of IgG2a/b antibodies and may have increased the efficacy of the vaccine by inducing a more balanced Th1/Th2 immune response, which was e.g. seen in the cytokine measurements from the splenocyte cultures of mice receiving the immunization with poly(I:C) and irradiated L3 larvae. Enhanced parasite-specific antibody levels and differences in isotype switching may also explain that the serum of mice immunized with a combination therapy performed superior in inhibiting larval motility in the ADCC *in vitro* assay. *In vivo*, the immunization with irradiated L3 larvae plus poly(I:C) or 3pRNA further reduced the frequency of MF-positive mice in comparison to the conventional immunization with att. L3 larvae alone, which is of importance, as it limits the transmission of the infection. Most importantly, the implementation of the nucleic acid receptor agonists improved vaccination efficacy, as the reduction in the adult worm burden was most prominent in mice immunized with a combination therapy of att. L3 larvae and poly(I:C) or 3pRNA, reaching a reduction of 73% and 57%, respectively. A similar beneficial impact of nucleic acid receptor agonists on vaccine efficacy was shown for *Schistosoma*, where the activation of the nucleic acid receptors TLR7/8 and TLR9 enhanced the immunization against *Schistosoma japonicum* (47). Similar to our results, their study observed enhanced production of Th-1 associated cytokines like IFN-γ and the authors suggested that this contributes to a reduced immunomodulation by regulatory T cells. Overall, the presented results indicate that type I IFNs may be protective during filarial infection and the targeting of the nucleic acids receptors TLR3, RIG-I and MDA5 by their agonists poly(I:C) and 3pRNA can enhance vaccination efficacy by strengthening protective immune responses.

## Materials and Methods

### Animals and infection

Six-week-old female BALB/c mice, C57BL/6 mice and eight-week-old *Meriones unguiculatus* were purchased from Janvier Labs, Saint-Berthevin, France. Albino C57BL/6 IFN-β Luciferase reporter mice were provided by Prof. Dr. Eicke Latz (Institute of Innate Immunity, University Hospital Bonn) and TLR7^-/-^, TLR9^-/-^, MDA5^-/-^ as well as STING^-/-^ mice were provided by Prof. Dr. Gunther Hartmann (Institute of Clinical Chemistry and Clinical Pharmacology, University Hospital Bonn). All animals were housed in individually ventilated cages within the animal facility at the IMMIP, University Hospital Bonn, with unlimited access to food and water and a 12 hour day/night cycle. All animal experiments were performed according to the EU Directive 2010/63/EU and approved by the state authorities Landesamt für Natur, Umwelt und Verbraucherschutz, Recklinghausen, Germany (protocol number 81-02.04.2020.A090, 84-02.04.2014.A327, 81-02.05.40.18.057).

Mice and *Meriones unguiculatus* were naturally infected with *L. sigmodontis* via the exposure to *Ornithonyssus bacoti* mites containing the infective L3 larvae for 24 hours and subsequent exchange of bedding for the following 7 days, as previously described (48).

### Parasite recovery

Infected *M. unguiculatus* were euthanized 5 days after the infection with an overdose of isoflurane (AbbVie, Wiesbaden, Germany) and *L. sigmodontis* L3 larvae were isolated by pleural lavage with Advanced-RPMI medium (37°C) (Thermo Fisher Scientific, Waltham, MA, USA). Mice were euthanized at the indicated time points with an overdose of isoflurane (AbbVie, Wiesbaden, Germany) and larvae or adult worms were isolated by pleural lavage with cold PBS (Thermo Fisher Scientific, Waltham, MA, USA). The isolated worms were quantified and the gender of adult worms was determined.

For microfilariae quantification, 50 µL peripheral blood were drawn from the facial vein to EDTA tubes and incubated in 1 mL 1x red blood cell lysis buffer (Thermo Fisher Scientific Inc., Waltham, Massachusetts, USA) for 10 min at room temperature (RT). Microfilariae in the pellet (400x*g*, 5 min, RT) were counted under the microscope.

### Organ preparation

#### Blood

Blood was collected from the facial vein of animals and stored at RT. Clotted samples were centrifuged (6000x*g*, 5 min, RT) and the serum was stored at -20 °C until analysis.

Mice were euthanized by an overdose of isoflurane (Piramal Critical Care Deutschland GmbH, Hallbergmoos, Germany). All cell counts were determined with a CASY® Cell Counter (Schärfe System GmbH, Reutlingen, Germany) using a 150 µm capillary.

#### Skin

The skin (approx. 1cm^2^) of the injection area was isolated. Half of the skin was stored in 700 µL Trizol (QIAGEN, Hilden, Germany) at -20°C until RNA isolation. The other half was minced and incubated at 37 °C on a shaker (350 rpm) for 75 min in RPMI medium (Life technologies Corporation, Grand Island, NY, USA), supplemented with 10% FCS (PAN Biotech, Aidenbach, Germany), 1% Penicillin (10,000 units/mL)/ Streptomycin (10 mg/mL) (Life technologies Corporation, Grand Island, NY USA), 2 mM L-Glutamine (Life technologies Corporation, Grand Island, NY USA), 0.25 mg/mL Liberase TL (Hoffmann-La Roche Ltd., Basel, Switzerland), and 0.5 mg/mL DNase I (Thermo Fisher Scientific, Waltham, MA, USA). The reaction was stopped with RPMI medium supplemented with 10 mM EDTA (Carl Roth, Karlsruhe, Germany) and 2% FCS was added. Cells were passed through a 70 µm cell strainer (Miltenyi Biotec, Bergisch Gladbach, Germany), centrifuged at 400x*g* for 5 min at 4 °C and taken up in MACS buffer (PBS, 10% FCS (PAN Biotech, Aidenbach, Germany), 2 mM EDTA (Carl Roth, Karlsruhe, Germany)). 1 × 10^6^ cells were analyzed by flow cytometry.

#### Pleura

The pleural lavage was performed with ∼ 5 mL cold PBS. The first mL was collected and centrifuged at 400*xg*, 5 min at 4 °C. The obtained supernatant was stored at -20 °C for ELISA analysis, the cell pellet was pooled with the remaining lavage fraction. Red blood cell lysis was performed with RBC buffer according to manufacturer’s protocol (Thermo Fisher Scientific Inc., Waltham, MA, USA). Cells were washed with MACS buffer (PBS, 1% FCS (PAN Biotech, Aidenbach, Germany), 2 mM EDTA (Carl Roth, Karlsruhe, Germany)) and 1 × 10^6^ cells were analyzed by flow cytometry.

#### Spleen

Isolated spleens were perfused by 0.5 mg/mL collagenase VIII buffer (Roche, Basel, Switzerland), minced and incubated at 37 °C for 30 min on a shaker with 350 rpm. MACS buffer (PBS, 1% FCS (PAN Biotech, Aidenbach, Germany), 2 mM EDTA (Carl Roth, Karlsruhe, Germany)) was added and a single cell suspension was generated by passing the cells through a 70 µm metal strainer. The cells were centrifuged at 400x*g* for 5 min at 4 °C. RBC lysis was performed according to the manufacturer’s protocol (Thermo Fisher Scientific Inc., Waltham, MA, USA). Afterwards the cells were washed with MACS buffer (PBS, 1% FCS (PAN Biotech, Aidenbach, Germany), 2 mM EDTA (Carl Roth, Karlsruhe, Germany)) and taken up in RPMI medium (Life technologies Corporation, Grand Island, NY, USA) supplemented with 10% FCS (PAN Biotech, Aidenbach, Germany), 1% Penicillin (10,000 units/mL)/ Streptomycin (10 mg/mL) (Life technologies Corporation, Grand Island, NY, USA) and 2 mM L-Glutamine (Life technologies Corporation, Grand Island, NY, USA). 1 × 10^6^ cells were analyzed by flow cytometry. For ELISA analysis, 2 × 10^6^ cells/mL were plated and stimulated with 2.5 µg/mL Concanavalin A (ConA, Sigma-Aldrich, St. Louis, MO, USA) or 25 µg/mL crude *L. sigmodontis* adult worm extract (LsAg) for 72 h. The supernatant was stored at -20 °C until analysis.

LsAg was prepared as previously described (49). In short, freshly isolated adult worms were rinsed in sterile PBS before being mechanically homogenized under sterile conditions. Insoluble material was removed by centrifugation at 400*xg* for 10 min at 4°C. Protein concentrations of crude extracts were determined using the Advanced Protein Assay Cytoskeleton, Denver, USA).

### Agonist injection and immunization protocol

20 µg/mouse poly(I:C) (HMW) VacciGrade™ (Invivogen, San Diego, CA, USA) and 20 µg/mouse 3pRNA (synthesized by AG Hartmann, University Hospital Bonn, Germany) were formulated with *in vivo*-jetPEI® (Polyplus-transfection SA, New York City, NY, USA) according to the manufacturer’s protocol at an N/P ratio of 8.

Attenuation of third stage larvae (L3s) by irradiation was performed at the department of Radiation Oncology, University Hospital Bonn, Germany. Radiation was performed using a TrueBeam STx® (Varian Medical Systems Germany). The photon energy of the radiation source was 10 MeV with a dose rate of 24 Gy/min. L3s were irradiated with 450 Gy applied in consecutive fractions without break with 100 Gy per fraction in a tissue equivalent RW3-Plasticphantom, at a depth of 23mm (dose maximum).

25 irradiated L3s and agonists were injected in separate syringes with a volume of 50µL each, right after the other in the same injection area at the hind leg for skin analysis or in the neck for all other experiments.

In order to analyze the immediate impact of the injections, skin and blood samples were taken 4 hours after the subcutaneous injection at the hind leg.

To identify the potential of nucleic acids receptor agonist to kill *L. sigmodontis* L3 larvae, mice were subcutaneously injected with an agonist at the neck half a day before the subcutaneous infection with 40 *L. sigmodontis* L3 larvae in the same neck area. This was followed half a day later by an intravenous injection of the same agonist. The pleural cavity was analyzed 9 days after the subcutaneous infection.

In the immunization experiments mice were subcutaneously injected at the neck with 25 L3 larvae attenuated by 450 Gy radiation (‘att.’) with or without an agonist for three times in two-week intervals. Two weeks after the last immunization, blood was drawn and mice were naturally infected (day 0). Further, blood was drawn on days 50, 57 and 63 and *ex vivo* analysis was performed on day 63 after the infection.

### Determination of parasite-specific antibodies

Plates were coated with 20 µg/mL LsAg diluted in PBS overnight (o/n) at 4 °C. Plates were washed with 0.5% Tween® 20 (Merck KGaA, Darmstadt, Germany) in 1% PBS, pH 7.2 and blocked for 2 h at RT with 1% bovine serum albumin (BSA) (PAA Laboratories, Cölbe, Germany) in PBS. Sera from day 0, isolated from blood that was drawn after the immunization and prior to the challenge infection, were diluted 1:50, sera from following time points after challenge infection were diluted 1:1000 in 1% BSA/PBS and incubated o/n at 4 °C. For *L. sigmodontis*-specific antibody quantification no standard is available. Therefore, a sample was prepared from pooled serum and loaded onto each plate. Thereby differences in OD due to varying incubation times between plates were assessed and values were recalculated in order to compare data measured on different plates. Plates were washed and biotinylated murine IgE, IgG1 or IgG2a/b antibodies (BD Biosciences San Jose, CA, USA) at a dilution of 1:400 in 1% BSA/PBS were added and incubated for 2 h at RT on a shaker. After washing, streptavidin-HRP (Thermo Fisher Scientific Inc., Waltham, MA, USA) was added for 30 min at RT on a shaker. After washing, TMB (Thermo Fisher Scientific Inc., Waltham, MA, USA) was added. Upon coloration the reaction was stopped with 2 M H_2_SO_4_ (Carl Roth GmbH + Co. KG, Karlsruhe, Germany). Reading was performed at 450 nm and 570 nm wavelength using a SpectraMax190 (Molecular Devices, San Jose, CA, USA) with Soft Max Pro 7 software (Molecular Devices, San Jose, CA, USA).

### In vitro motility assay

The assay was adapted from Veerapathran *et al*. (50). L3 larvae were recovered by pleural lavage from *M. unguiculatus* 5 days after natural *L. sigmodontis* infection. Peritoneal cells were isolated from naïve BALB/c WT donor mice. 2 × 10^5^ peritoneal cells were co-cultured with 10-12 L3 larvae in RPMI-medium (Life technologies Corporation, Grand Island, NY, USA) supplemented with 25% pooled serum drawn from immunized animals prior to challenge infection (two weeks after the final vaccination). The motility of L3 larvae was scored under the microscope on a daily basis for a total of three days. Following score was used to assess the motility: 4: fast and continuous movement, 3: slower but continuous movement, 2: slower and discontinuous movement, 1: movement discontinuous and limited to larval ends, 0: no movement observed within 30 seconds.

### Cytokine quantification by ELISA

Cytokines were quantified in the first mL of pleural wash and in the 72 h splenocyte culture supernatant. IL-5 and IFN-γ were quantified using Invitrogen™ eBioscience™ ELISA Ready-SET-Go!™ (Thermo Fisher Scientific, Waltham, MA, USA). IP-10 (CXCL10), RANTES (CCL5) and Eotaxin 1 (CCL11) were quantified using DuoSet ELISA kits (R&D Systems, Minneapolis, MN, USA). The manufacturers’ protocol was followed.

2 M H_2_SO_4_ (Carl Roth GmbH + Co. KG, Karlsruhe, Germany) served as stopping solution. Reading was performed at 450 nm and 570 nm wavelength using a SpectraMax190 (Molecular Devices, San Jose, CA, USA) with Soft Max Pro 7 software (Molecular Devices, San Jose, CA, USA).

### Flow cytometric analysis of skin, pleura and spleen cells

1 × 10^6^ cells were pelleted (400x*g*, 5 min, 4°C). For surface staining cells were incubated with mastermix prepared in Fc-block (1%FCS/PBS with 0.1% rat IgG (Sigma-Aldrich, St. Louis, MO, USA)). Master mixes were prepared as combinations of following antibodies. If not stated otherwise the antibodies were purchased from BioLegend, San Diego, CA, USA: CD11b (Al700, clone M1/70), CD11c (BV605, clone N418), CD45 (PerCPCy5.5, clone 30-F11), CD86 (Al647, clone GL-1), I-Ab (PE-Cy7, clone AF6-120.1), Ly6C (APC-Cy7, clone HK1.4), Ly6G (BV421, clone 1A8), purified RELM-α polyclonal, rabbit, (PeproTech, Inc., Rocky Hill, NJ, USA) combined with goat anti-rabbit Al488 (Invitrogen, Carlsbad, CA, USA), SiglecF (PE, clone E50-2440, BD, San Jose, USA). For intracellular staining cells were incubated in fixation/permeabilization buffer (Thermo Fisher Scientific Inc., Waltham, Massachusetts, USA) for 20 min at RT. Cells were washed and blocked overnight in Fc-block (1 % bovine serum albumin fraction V (BSA) (PAA Laboratories, Cölbe, Germany) in PBS, 1:1000 rat IgG (Sigma-Aldrich, St. Louis, MO, USA)) at 4°C. The next day, cells were permeabilized with permeabilization buffer for 20 min at RT (Thermo Fisher Scientific Inc., Waltham, Massachusetts, USA) and stained with mastermix containing antibodies for extracellular and intracellular targets for 45 min at 4 °C. After staining cells were washed. Data acquisition was performed on a CytoFLEX S (Beckman Coulter, Brea, CA, USA) and analysis with FlowJo® Software V10 (FlowJo, LLC, Ashland, OR, USA). Fluorescence minus one (FMO) controls were used for evaluation.

### Luciferase IFN-β reporter assay

2 × 10^6^ pleura cells were plated and stimulated with 1 µg/mL poly(I:C) (Invivogen, San Diego, CA, USA) for 4 h at 37 °C. Cells were lysed in a 96-well flat-bottom white plate (Thermo Fisher Scientific Inc., Waltham, Massachusetts, USA) using 50 µL Glo lysis buffer (Promega Corporation, Madison, Wisconsin, USA) for 5 min at RT on a shaker. 50 µL of ONE-Glo™ luciferase assay substrate (Promega Corporation, Madison, Wisconsin, USA) was added and cells were incubated for 10 min at RT on a shaker. Luminescence was measured with an Infinite® M-plex (Tecan Trading AG, Männedorf, Switzerland) and i-control™ software (Tecan Trading AG, Männedorf, Switzerland) with an integration time of 1 second.

### RNA isolation

Skin samples stored in 700 µL Trizol were homogenized in a Precellys® 2 mL Soft Tissue Homogenizing Ceramic Beads Tube (Cayman Chemical, Ann Arbor, MI, USA) using the Precellys® 24 machine, program “6000-3×60-120”. The homogenate was incubated at RT for 5 min in a fresh vial. 70 µL 1-Bromo-3-chloropropane (BCP) (Tokyo Chemical Industry, Tokyo, Japan) were added, the sample was vortexed and incubated for 2-3 min at RT. After centrifugation for 15 min at 12000x*g* at 4 °C, 350 µL of aqueous phase was transferred into a 2 mL reaction tube and subjected to the QIAcube (QIAGEN, Hilden, Germany) for RNA isolation. The animal tissue and cell protocol including an on-column DNase digest was followed using the RNeasy® Mini kit (QIAGEN, Hilden, Germany). The 70% ethanol was exchanged with 100% ethanol.

### cDNA and RT-PCR

cDNA was generated from 1 µg RNA with the Omniscript® Reverse Transcription Kit (Qiagen, Hilde, Germany) using oligoDT_12-18_ primer (Thermo Fisher Scientific Inc., Waltham, Massachusetts, USA) and RNaseOUT™ recombinant ribonuclease inhibitor (Thermo Fisher Scientific Inc., Waltham, Massachusetts, USA). The mastermixes were prepared using the HotStarTaq® DNA Polymerase kit (QIAGEN, Hilden, Germany) and SYBR™ Green Nucleic Acid Stain (Thermo Fisher Scientific Inc., Waltham, MA, USA). Samples were run on a Rotor-Gene Q (QIAGEN, Hilden, Germany) and analyzed with Rotor-Gene Q Series software (QIAGEN, Hilden, Germany).

Primer sequences: β-actin: forward: 5’ TGACAGGATGCAGAAGGAGA 3’, reverse: 5’ CGCTCAGGAGGAGCAATG 3’. IP-10/CXCL10: forward: 5’ GCCGTCATTTTCTGCCTCAT 3’, reverse: 5’ GCTTCCCTATGGCCCTCATT 3’. IFN-β: forward: 5’ CAGGCAACCTTTAAGCATCAG 3’, reverse: 5’ CCTTTGACCTTTCAAATGCAG 3’.

### Statistical analysis

GraphPad Prism software version 8 (GraphPad Software, San Diego, CA, USA) was used for statistical analysis. Kruskal-Wallis test followed by Dunn’s post hoc multiple comparisons were used to test for significant differences between multiple groups. Mann-Whitney-U-test was used to test for significant differences between two unpaired groups. Data are shown as median with interquartile ranges. p values <0.05 were considered as significant. Prior to pooling data from different experiments, data were analyzed for homogeneity by not passing Spearman’s test for heteroscedasticity. If data could not be pooled, but statistical trends (p<0.1) were confirmed by repeated experiments, the trends were indicated in the figure.

## Acknowledgements

Images for experimental setups shown in Fig. 2A, 3A, 4A and 4E were taken from Servier Medical Art by Servier, which is licensed under a Creative Commons Attribution 3.0 Unported License.

## Data Availability Statement

The original contributions presented in the study are included in the article/supplementary material, further inquiries can be directed to the corresponding author/s.

## Author contributions

Conceptualization: JFS, CC, BS, AH, MPH.

Data curation: JFS, CC, BS, MPH.

Experimentation: JFS, FR, JJR, BL, SG, SJF, ALN, MK.

Investigation: JFS, FR, JJR, BL, SG, AN, SG, MK

Resources: SG, MR, EL, CC, GH, AH, MPH.

Supervision: CC, BS, AH, MPH.

Writing - original draft: JFS, JA, MPH.

Writing - review & editing: all authors.

## Supporting information

**S1 Table: Prescreening of potential adjuvants**.

**S1 Figure: Changes in pleural cell composition after treatment with poly(I:C) or 3pRNA**.

**S2 Figure: Gating strategy of peritoneal exudate cells**.

**S3 Figure: Immunization-induced worm clearance affects female and male *L. sigmodontis* filariae similarly**.

